# Genetic targeting of myelinated primary afferent neurons using a new *Nefh^CreERT2^* knock-in mouse

**DOI:** 10.1101/2025.03.18.643951

**Authors:** John CY Chen, Lech Kaczmarczyk, Felipe Meira de Faria, Marcin Szczot, Walker S Jackson, Max Larsson

**Affiliations:** Department of Biomedical and Clinical Sciences, Division of Cell- and Neurobiology, Linköping University, S-581 85 Linköping, Sweden; Department of Biomedical and Clinical Sciences, Center for Social and Affective Neuroscience, Linköping University, S-581 85 Linköping, Sweden

## Abstract

Primary afferent neurons that convey somatosensory modalities comprise two large, heterogeneous populations: small-diameter neurons that give rise to slowly conducting unmyelinated axonal C fibers and medium-to-large diameter neurons with fast myelinated A fibers. Despite these two major groupings, tools to differentiate between unmyelinated and myelinated primary afferent fibers by genetic targeting have not been available; in particular, whereas numerous mouse driver lines exist to target different C fiber populations, genetic tools that target myelinated primary afferent populations are scarce. Here we describe a knock-in mouse line expressing tamoxifen-dependent CreERT2 under control of the *Nefh* gene, which encodes neurofilament heavy chain (NFH or NF200), a protein that is highly enriched in myelinated fibers. This mouse enables highly selective and efficient recombination of Cre-dependent reporters for functional and anatomical interrogation of myelinated fibers while excluding unmyelinated C fibers. In combination with other recombinase-expressing mouse lines, this genetic tool will be valuable for intersectional targeting of subpopulations of myelinated primary afferent fibers.

## Introduction

Somatosensory stimuli are detected by primary afferent nerve fibers (PAFs), the cell bodies of which reside in the dorsal root ganglia (DRGs). A long-standing effort within the somatosensory research field has been to classify different populations of DRG neurons according to which somatosensory modalities they subserve. This has culminated in recent years with numerous single cell RNA sequencing studies identifying distinct clusters in a variety of species [e.g., 1,2–5]. However, functional and anatomical characterization of such clusters (and possible subclusters) has in many cases been hampered by a lack of unique genetic markers that could be used for targeting. Moreover, similar gene expression in other tissues, especially in other parts of the nervous system [6], may also complicate genetic manipulation based on a single marker. One potential resolution to this issue may be to employ intersectional targeting that relies on two or more genetic markers that are more likely to be uniquely expressed in the population of interest.

The earliest scheme for classification of somatosensory nerve fibers was based on the work by Erlanger and Gasser, who noted that axonal diameter and degree of myelination correlate with conduction velocity and that fibers could be grouped according to these properties [7,8]. Unmyelinated fibers, called C fibers, have small axonal diameter and conduct at very low velocities in the range of 0.5-1 m/s. Myelinated fibers are divided into thickly myelinated, thick-diameter Aα/β fibers with the highest conduction velocities, and thinly myelinated, smaller-diameter Aδ fibers with intermediate conduction velocities. The Erlanger-Gasser classification was found to be associated with physiological function, in that Aα fibers are proprioceptors, Aβ fibers generally low-threshold mechanoreceptors (LTMRs), Aδ fibers cool receptors (in humans) or fast nociceptors, and C fibers warm receptors, slow nociceptors or itch receptors [8]. While this relation between myelination and function has long been known to not be absolute, with the identification of C fiber LTMRs (C-LTMRs) [9–11] and ultrafast Aβ nociceptors [12,13], myelination remains a major distinguishing feature for different populations of PAFs.

One of the earliest protein markers identified that distinguishes separate populations of PAFs was the heavy chain of neurofilament protein, NFH (also called NF200; encoded by the *Nefh* gene), which was found to be selectively expressed in myelinated neurons [14,15]. Specifically, the expression of NFH among such neurons showed a positive correlation with conduction velocity, such that thickly myelinated, large-diameter Aα/β neurons on average exhibit the highest levels of NFH, while thinly myelinated, smaller-diameter Aδ neurons express the protein at lower levels [15]. By contrast, unmyelinated small-diameter C fibers show little or no NFH expression. This observation has been corroborated in recent transcriptomic studies, showing that whereas C fibers express *Nefh* only very weakly, likely Aβ fibers exhibit high expression, and putative Aδ fibers have intermediate levels of expression of *Nefh* [1,3].

Thus, *Nefh* could be a useful genetic marker for selective targeting of myelinated PAFs, and in intersectional targeting strategies where a second marker is expressed in both myelinated and unmyelinated PAFs, or in other cells that do not express *Nefh*. To that end, we have in the present study generated a knock-in mouse line expressing tamoxifen-inducible CreERT2 recombinase from the endogenous *Nefh* locus. We show this tool to be a highly efficient tool for selective and specific genetic manipulation of myelinated PAFs.

## Materials and methods

### Animals

The *Nefh^CreERT2^* knock-in mouse line was generated using **a standard** CRISPR/Cas9 **approach** on a C57Bl/6J background (Fig. 1A). To construct the targeting construct, homology arms were captured by PCR amplification from C57Bl/6J genomic DNA. An IRES2 sequence (encephalomyocarditis virus) was inserted 55 bp downstream of the translation termination codon (TAA), and immediately followed by coding sequence for Cre-ERT2 obtained from a plasmid we used previously [16]. **The gene editing procedure was performed** within the Laboratory of RNA Biology and Mouse Genome Engineering Facility **(IN-MOL-CELL Infrastructure)** at International Institute of Molecular and Cell Biology in Warsaw, Poland. **In brief, C57Bl/6J zygotes obtained from mated, hormonally stimulated females were microinjected into the cytoplasm using Eppendorf 5242 microinjector (EppendorfNetheler-Hinz GmbH) and Eppendorf Femtotips II capillaries with the following CRISPR cocktail: Cas9 mRNA (25 ng/µl), sgRNA (15 ng/µl), and donor dsDNA (7.5 ng/µl). After overnight culture microinjected embryos at 2-cell stage were transferred into the oviducts of 0.5-day p.c. pseudo-pregnant females. Born pups were genotyped by PCR at around 4 weeks. The presence of mutation was confirmed by Sanger sequencing in the founder mouse and, after backcrossing, in N1 generation mice. Routine g**enotyping was performed using the following primers: wt forward, 5’-CAG CGG CAC CAG AGA AGA AAG AC-3’; mutant forward, 5’-AGA ACG TGG TGC CCC TCT ATG ACC-3’; common reverse, 5’-CAG GCC CAC CAT CTA AGC AGT GT-3’. PCR reactions were performed using 30 cycles with annealing at 60°C for 30 s, and extension at 72°C for 20 s. These reactions yield products of 270 bp for the Cre allele and 358 bp for the wild-type allele and can identify wild-type, homozygous and heterozygous **allele configurations**.

**Figure 1.**
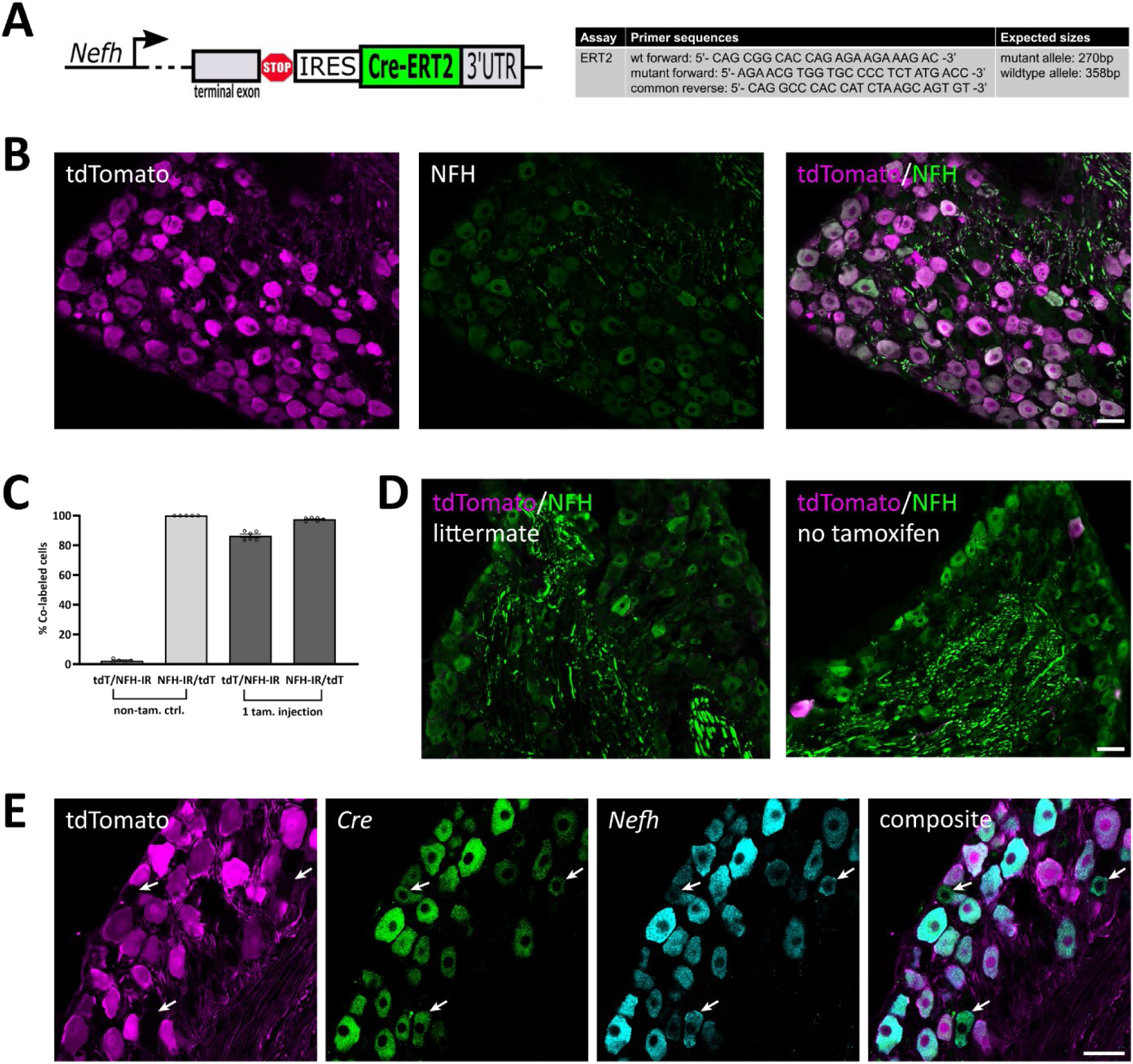
Verification of the *Nefh*^CreERT2^ mouse line in DRGs. **A**, Design of the allele and genotyping primers. **B**, co-localization of tdTomato and NFH immunoreactivity (IR) in a lumbar DRG after a single injection of tamoxifen (2 mg) in the adult. **C**, quantification of the co-localization of tdTomato and NFH-IR after one tamoxifen injection or in non-injected mice (mean ± S.E.M.). Each data point indicates a section from a ganglion; n=3 mice. **D**, example images of tdTomato expression in littermate Cre-negative mice and in Cre expressing mice that had not been subjected to tamoxifen injection. **E**, co-localization of tdTomato with *Cre* and *Nefh* mRNA transcripts. Arrows indicate *Nefh*^+^ cells that do not show tdTomato fluorescence but are *Cre*^+^. Scale bars are 50 µm for all panels.

For the purposes of characterization of the *Nefh^CreERT2^* mouse line, mice from this line were crossed with mice from the following lines: RCL-tdTomato (Ai14; Jackson Laboratory, strain #007914) [17]; RCL-GCaMP6f (Ai95D, Jackson Laboratory, strain #028865) [18], and LSL-APEX2 mice. The latter strain was derived by crossing ROSA26^DR-Matrix-dAPEX2^ (Jackson Laboratory, strain #032764), which expresses dAPEX2 peroxidase localized to the mitochondrial matrix in a Cre- and Flp-dependent manner [19], to ROSA26Flpo mice (Jackson Laboratory, strain #012930) [20] for deletion of the Frt-stop-Frt cassette. Animal experiments are reported according to the ARRIVE guidelines. All animal experiments were approved by the Animal Ethics Committee at Linköping University (permit no. 2439-2021) and performed in accordance with the EU Directive 2010/63/EU.

### Tamoxifen administration

Tamoxifen (Sigma-Aldrich T5648) was dissolved in corn oil at 20 mg/mL (or, in three animals, 0.2 mg/mL) concentration overnight at 37°C. Mice at 6-10 weeks of age received either one, two or five tamoxifen injections (100 µL, i.p.). For two injections, these were administered one day apart while in the case of five injections they were administered once daily. The mice were subjected to further experiments not earlier than two weeks after tamoxifen injections.

### Tissue preparation

Mice of either sex were anesthetized with sodium pentobarbital (100 mg/kg) and subjected to transcardial perfusion using 5 mL phosphate buffer (PB, 0.1 M, pH 7.4) followed by 50 mL of 4 % paraformaldehyde, or (for electron microscopy) either 2 % paraformaldehyde and 2.5 % glutaraldehyde or 4 % paraformaldehyde and 0.5 % glutaraldehyde. Tissue (brain, spinal cord, dorsal root ganglia, sciatic nerves, back skin and hind paw skin) was harvested, post-fixed in fixative at 4°C overnight and stored in 1/10 fixative at 4°C before further processing.

### Immunofluorescence

Tissue specimens from *Nefh*^CreERT2/wt^;Ai14^+/wt^ mice (12 females, 6 males, 9-12 weeks old) and *Nefh*^wt/wt^;Ai14^+/wt^ littermate controls (2 females, 9 weeks) were cryoprotected in 30 % sucrose and embedded in OCT. Sections were cut at 15-20 µm thickness in a cryostat and placed on slides. The sections were incubated in phosphate-buffered saline (PBS) with 3% normal goat serum, 0.5 % bovine serum albumin and 0.5 % Triton X-100 (blocking solution), followed by incubation in primary antibodies (see Table 1) diluted in blocking solution at room temperature overnight. After rinsing, the sections were incubated in blocking solution containing secondary antibodies at a dilution of 1:500. In cases where two primary antibodies raised in rabbit were used, a tyramide signal amplification (TSA) approach was used to differentiate between them. In short, the sections were sequentially incubated in the primary antibody against tdTomato diluted at 1:10 000, biotinylated goat anti-rabbit (dilution 1:200), streptavidin-horseradish peroxidase (1:100; Life Technologies), and tyramide-Alexa 568 (Life Technologies); after this, the labelling procedure continued with the second primary antibody. To detect binding sites for isolectin B_4_ (IB_4_), biotinylated IB_4_ (dilution 1:2000) in conjunction with streptavidin-Alexa 488 (dilution 1:500; Life Technologies) was used. In skin sections, cellular nuclei were labeled in a separate step using DAPI (1:1000; Life Technologies) or SYTOX Deep Red (1:2000). Sections were coverslipped with Prolong Diamond or SlowFade Diamond (Life Technologies). Sections were imaged using a Zeiss LSM800 confocal microscope, or a Leica DMi8 widefield microscope.

**Table 1.**
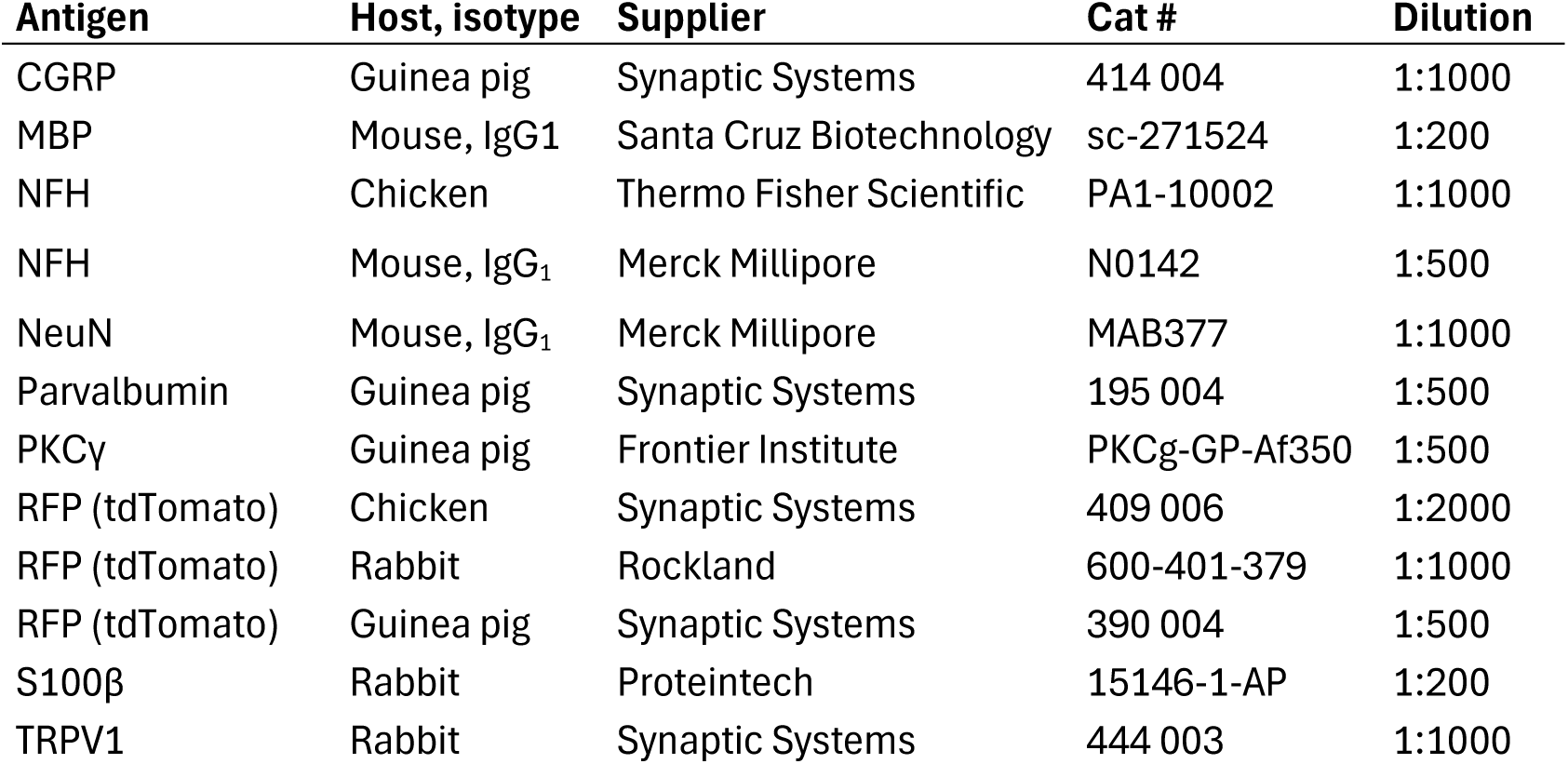
Primary antibodies.

To determine recombination efficiencies and co-localization with cell or axonal markers, L3-L5 DRGs from *Nefh*^CreERT2/wt^;Ai14^+/wt^ mice subjected to different tamoxifen regimes were analyzed: no tamoxifen injection, 2 animals (females, 12 weeks); single injection of low-dose (20 µg) tamoxifen, three animals (females, 11 weeks); single injection of normal dose (2 mg) tamoxifen, three animals (females, 10 weeks); two injections of normal dose tamoxifen, 2 animals (females, 11 weeks); five injections of normal dose tamoxifen, 3 animals (males, 10 weeks). Two *Nefh*^wt/wt^;Ai14^+/wt^ littermate controls (females, 9 weeks) received a single normal dose tamoxifen injection. Only normal tamoxifen dose/single injection animals were used for quantification of co-localization with cell/axonal markers. No randomization or blinding with respect to genotype or tamoxifen regime was performed. Additional *Nefh*^CreERT2/wt^;Ai14^+/wt^ mice (3 females, 2 males, 9-11 weeks) subjected to single injection of normal tamoxifen dose were used for qualitative morphological and immunohistochemical analysis.

### In situ hybridization

The protocol for *in situ* hybridization chain reaction (HCR) is based on the Molecular Instruments’ HCR RNA-FISH protocol for frozen fixed tissue sections. In brief, DRG sections of 15 µm thickness were incubated in probe hybridization buffer with 0.4 pmol of each probe set overnight at 37°C. Excess probes were then removed by incubation of probe wash buffer and 5X SSCT (sodium chloride sodium citrate with 0.1% Tween 20). Before amplification, 6 pmol of hairpin h1 and 6 pmol of hairpin h2 were subjected separately to snap cooling (95°C for 90 s, followed by cooling at room temperature in a dark drawer for 30 min). During amplification stage, the sections were incubated in amplication buffer with h1 and h2 hairpins overnight at room temperature. Excess hairpins were then removed by incubation of 5X SSCT. Sections were coverslipped with Prolong Diamond and imaged using a Zeiss LSM800 confocal microscope.

### Image analysis

For measuring the soma size of DRG neurons, z-stacks (12-16 images) of each DRG section were obtained using a Zeiss LSM800 confocal microscope. The cross-sectional area of each DRG neuron was determined by identifying the cross section that was larger than both the previous and the next images in the z-stack. The cross-sectional areas were then measured using the Fiji software.

### Electron microscopy

Spinal cord from *Nefh*^CreERT2/wt^;LSL-APEX2^+/wt^ mice (1 female, 3 males, 14 weeks old) and one female *Nefh*^wt/wt^;LSL-APEX2^+/wt^ littermate control (14 weeks) was embedded in 4 % low-melting agarose (Fisher Scientific #10377033) and sectioned on a vibrating microtome (Campden Instruments 7000smz-2) into transverse 150 µm sections. For APEX2 histochemistry, sections were pre-incubated in 3,3′-diaminobenzidine (DAB; Vector Laboratories) without H_2_O_2_ for 30 min, and then incubated in DAB with H_2_O_2_ for 90 min. The tissue was then osmicated in 1 % OsO_2_, dehydrated in a graded series of ethanol and embedded in Durcupan ACM (Sigma-Aldrich #44610). Ultrathin sections (70 nm thickness) were cut and placed on single-slot copper grids. Sections were counterstained using 2 % uranyl acetate and 0.5 % lead citrate before examination in a JEOL JEM1400 Flash transmission electron microscope at 80kV.

### *In vivo* imaging

*Nefh*^CreERT2^ animals were crossed with RCL-GCaMP6f mice to generate *Nefh*^CreERT2/wt^;GCaMP6f^+/wt^ mice. In control experiments pan-neuronal GCaMP6f expression was obtained using postnatal (P0-P2) i.p. injection of AAV9-CAG-Cre into RCL-GCaMP6f pups [pan-GCaMP6f; 21]. For *Nefh*^CreERT2^;GCaMP6f mice, tamoxifen was administered as above. Mice of both sexes (*Nefh*^CreERT2^;GCaMP6f: **2 males**, 4 females; pan-GCaMP6f: 2 females, 2 males) **9-16** weeks old were used.

For imaging, the mouse was anesthetized with isoflurane (4% induction, 1,5% maintenance) and transferred to a custom surgical platform equipped with a heating pad to maintain body temperature. The dorsal aspect of the spinal cord was surgically exposed from L5 to S2 vertebrae and stabilized with a spinal clamp (Narishige STS-A). To get optical access to the DRGs L6 and S1, a dental drill was used to chip the bone surface, including the spine of each targeted vertebra towards both the articular and the lateral process. Following surgery, the animal was transferred to the stage of a custom tilting light microscope (Thorlabs Cerna) equipped with a 4x, 0.28 NA air objective (Thorlabs). GCaMP6f fluorescence images were acquired for 40 second epochs at 5 Hz with an sCMOS camera (Sona 4.2, Andor) using a standard green fluorescent protein (GFP) filter cube. Stimuli were applied to the the hairy skin around the anogenital area of the mouse. To verify which stimuli recombined cells responded to, we applied a set of stimuli **in *Nefh*^CreERT2^;GCaMP6f mice (n=2)** consisting of air puff, vibration (100 Hz), gentle brushing, hair pull, heating (22°C → 48°C → 22°C) and cooling (22°C → 8°C). In another experimental set up we compared the proportions of responding cells in pan-GCaMP6f and *Nefh*^CreERT2^;GCaMP6f mice. In this setup the stimuli were gentle brush, hair pull, and pinch. Analysis of calcium imaging was performed as previously described [22]. Regions of interest (ROI) outlining responding cells were drawn in FIJI/ImageJ **using the Cell Wand Tool plugin** and relative change of GCaMP6f fluorescence was calculated as percent ΔF/F. Contaminant signal, *e.g*., from out-of-focus tissue and neighboring cells, was removed by subtracting the fluorescence of a donut-shaped area surrounding each ROI using a custom MATLAB script.

Overlapping ROIs and rare spontaneously active cells were not included in the analysis; threshold was set at 20 %. Imaging episodes were concatenated for display as traces or activity heatmaps where final ΔF/F was **shown** from 20 to 100%. Spatial maps of activity were generated by calculating the standard deviation for each pixel over a stimulation episode in FIJI/ImageJ.

### Statistical analysis

GraphPad Prism was used for descriptive statistics and Fisher’s exact test.

## Results

Mice hetero- or homozygous for the *Nefh^CreERT2^* allele were viable and fertile, and did not exhibit any overt behavioral phenotype either when bred alone or when crossed with any of the reporter lines in this study. To characterize recombination in these mice, *Nefh^CreERT2^*; Ai14 mice were injected with tamoxifen at 6-10 weeks of age. Because expression of *Nefh* transcript and NFH protein during embryonic and early postnatal development is very low [3,23], we did not examine mice where tamoxifen was administered at an earlier age. In lumbar (L3-L5) DRGs from mice injected once with 2 mg tamoxifen, 98 % of neurons with endogenous tdTomato fluorescence were also immunoreactive (IR) for NFH, whereas 86 % of NFH-IR neurons were tdTomato^+^ (Fig. 1B, C). In littermate *Nefh^CreERT2wt/wt^*;Ai14^+/wt^ mice DRG neurons were never tdTomato^+^, and no or very few neurons exhibited tdTomato fluorescence in *Nefh^CreERT2+/wt^*; Ai14^+/wt^ mice that had not received tamoxifen (Fig. 1C, D). *In situ* hybridization further showed essentially complete co-localization between *Cre* and *Nefh* mRNA in lumbar DRGs (Fig 1E).

Among DRG neurons, myelination shows a positive correlation with soma size, such that myelinated primary afferents arise from medium-to-large-sized neurons whereas unmyelinated neurons are generally small [15]. We therefore analyzed the cross-sectional area of DRG neurons in L3-5 ganglia that showed recombination in the *Nefh^CreERT2^*;Ai14 mice. In mice receiving one 2 mg tamoxifen injection, tdTomato^+^/NeuN-IR neurons were generally medium-to-large, whereas tdTomato^-^/NeuN-IR neurons were mostly small (Fig. S1A). The same size distributions were found in mice receiving a single low-dose (20 µg) tamoxifen injection, or two or five injections of 2 mg tamoxifen (Fig. S1B-D), although there was a tendency towards a larger proportion of small neurons being labelled with more tamoxifen injections. In subsequent experiments, one 2 mg tamoxifen injection was used to induce recombination. To further corroborate that recombination occurred in neurons giving rise to myelinated PAFs, sciatic nerve sections from *Nefh^CreERT2^*;Ai14 mice were immunolabelled for NFH and myelin basic protein (MBP), which labels the myelin sheath (Fig. 2). Here, tdTomato^+^ axons showed near-complete co-localization with both NFH-IR and MBP-IR. Because the sciatic nerve is a mixed sensory/motor nerve, this also indicates tdTomato^+^ expression in the myelinated axons from spinal motor neurons. In the skin, fibers were generally restricted to the dermis, although some fibers penetrated into the epidermis (Fig. 3). tdTomato^+^ fibers associated with S100β^+^ Meissner corpuscles in dermal papillae were observed, as were tdTomato^+^ circumferential and lanceolate endings around hair follicles. Combined, these observations indicate that the Nefh^CreERT2^ mouse line robustly targets *Nefh* expressing, myelinated DRG neurons with very high specificity and selectivity.

**Figure 2.**
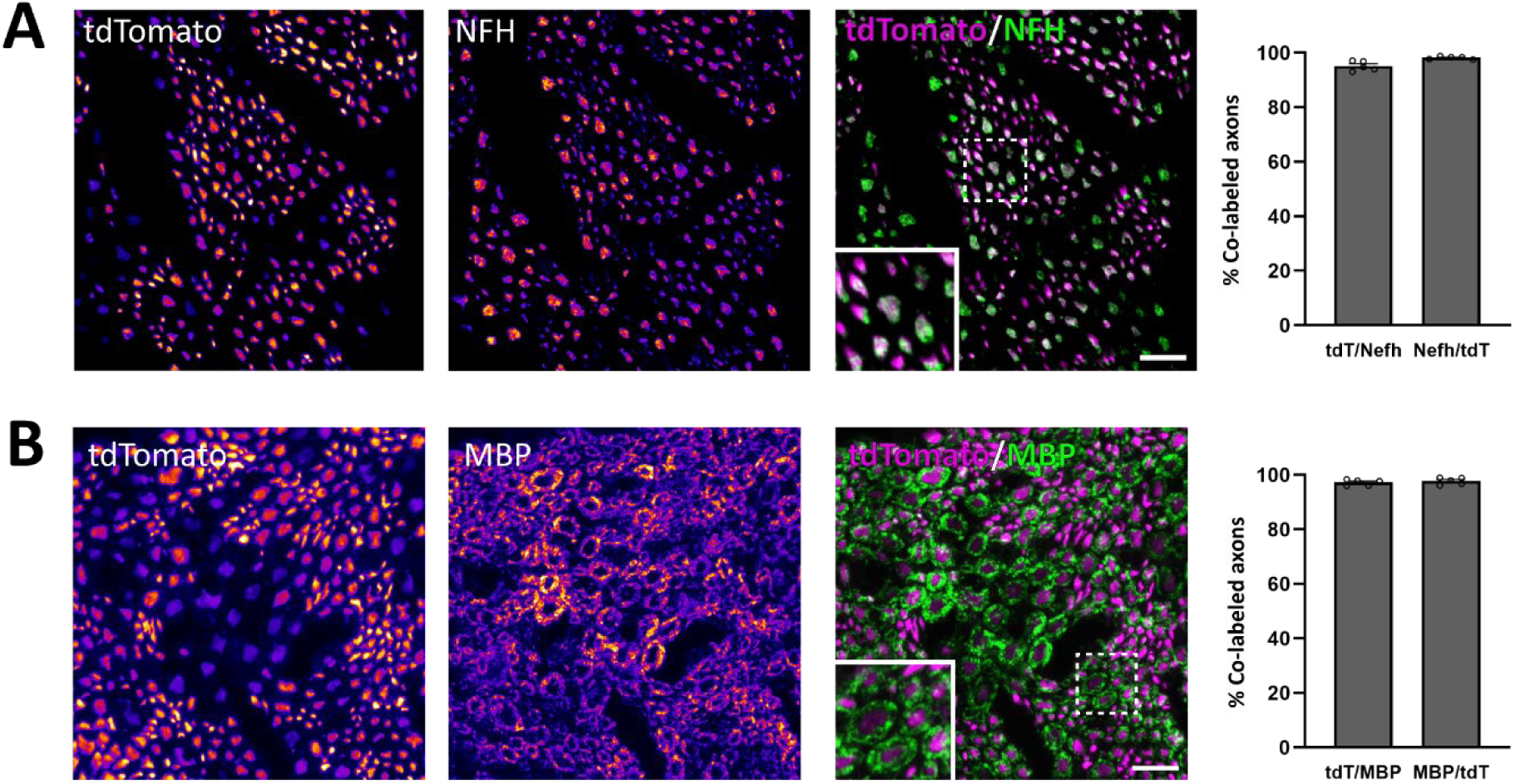
Reporter expression in peripheral nerve axons. **A**, Colocalization of tdTomato and NFH in axons in the sciatic nerve. **B**, tdTomato expression in axons enveloped by MBP-IR myelin sheaths. The two left-most panels show immunoreactivity for tdTomato and NFH/MBP**, pseudocolored using the “Fire” look-up table in FIJI to increase visibility of weak immunoreactivity**. **Insets show the regions indicated by dashed frames at higher magnification.** Data are presented as mean ± S.E.M.; n=3 mice. Scale bars, 10 µm.

**Figure 3.**
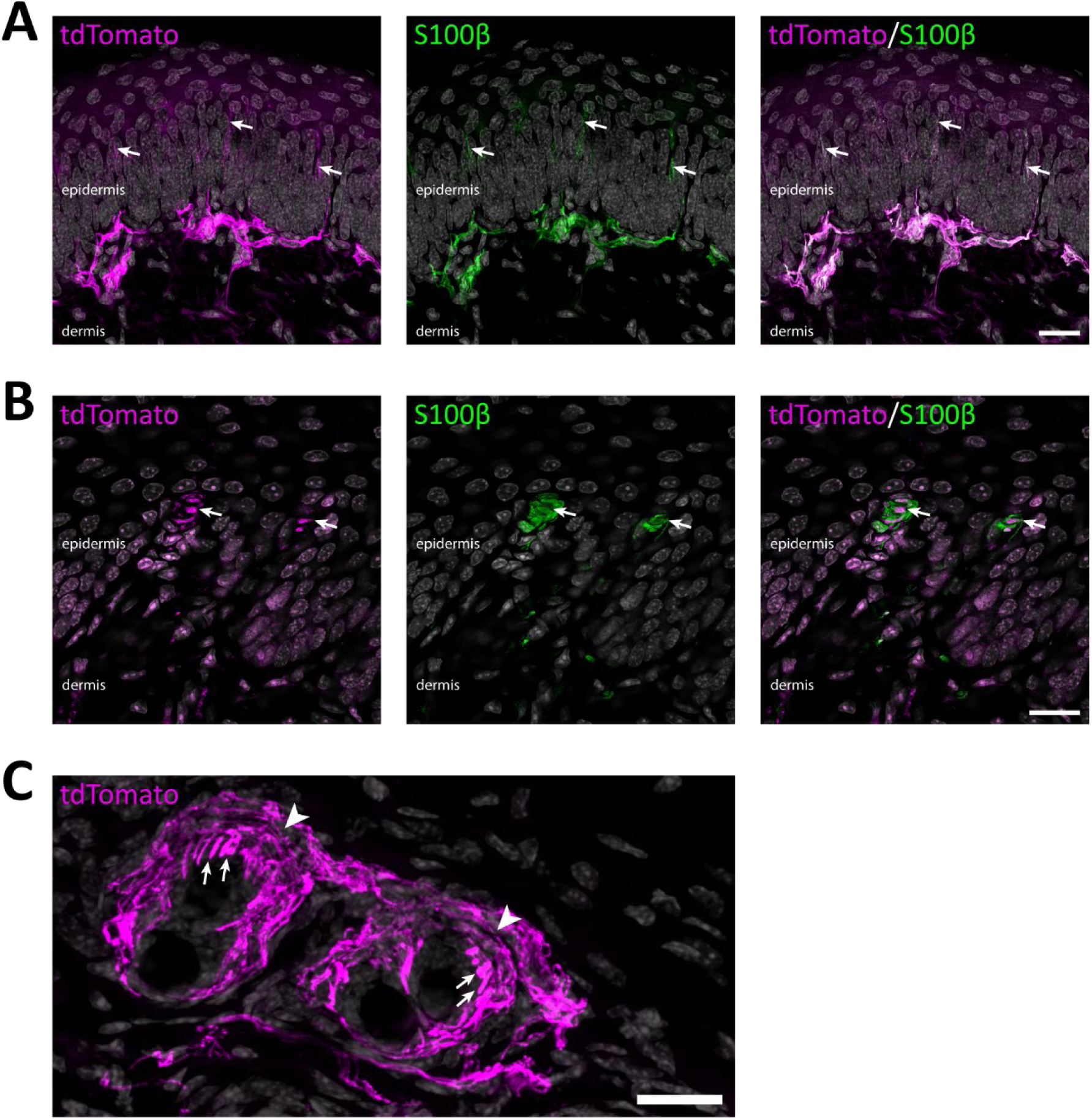
Reporter-expressing fibers in skin. **A**, Co-localization of tdTomato^+^ fibers with S100β, a marker for myelinated axons and myelinating Schwann cells, in hind paw glabrous skin. Most fibers are restricted to the dermis, but some protrude into the epidermal layer (*arrows*). Maximum intensity projection of 17 deconvolved optical sections at 0.5 µm separation. **B**, tdTomato^+^ fibers innervating S100β-IR Meissner corpuscles (*arrows*) in hind paw glabrous skin. A single deconvolved optical section. **C**, an oblique top view of a hairy skin section. tdTomato^+^ fibers formed circumferential (arrowheads) and lanceolate (arrows) endings around hair follicles. Maximum intensity projection of 30 optical sections at 0.53 µm separation. In all panels, cell nuclei are labeled by DAPI (grey). Scale bars, 20 µm.

To further characterize DRG populations targeted in the *Nefh*^CreERT2^ mouse, L3-5 DRG sections from *Nefh*^CreERT2^;Ai14 mice were labelled for a panel of common markers of different DRG neuron populations (Fig. 4). IR for PV, a marker of proprioceptors [24], was found in 53 ± 3 % (mean ± S.E.M.) of tdTomato^+^ neurons, whereas 85 ± 2 % of PV-IR neurons were tdTomato^+^. As expected, there was no overlap with binding of isolectin B_4_ (IB_4_), which marks a population of largely non-peptidergic C fibers that show no NFH-IR [8,25]. By contrast, CGRP, a marker for peptidergic C and A fibers, was found in 26 ± 2 % of tdTomato^+^ neurons, and 51 ± 4 % of CGRP-IR neurons were tdTomato^+^; TRPV1-IR was found in 10 ± 1 % of tdTomato^+^ neurons, while 13 ± 1 % of TRPV1-IR neurons were tdTomato^+^.

**Figure 4.**
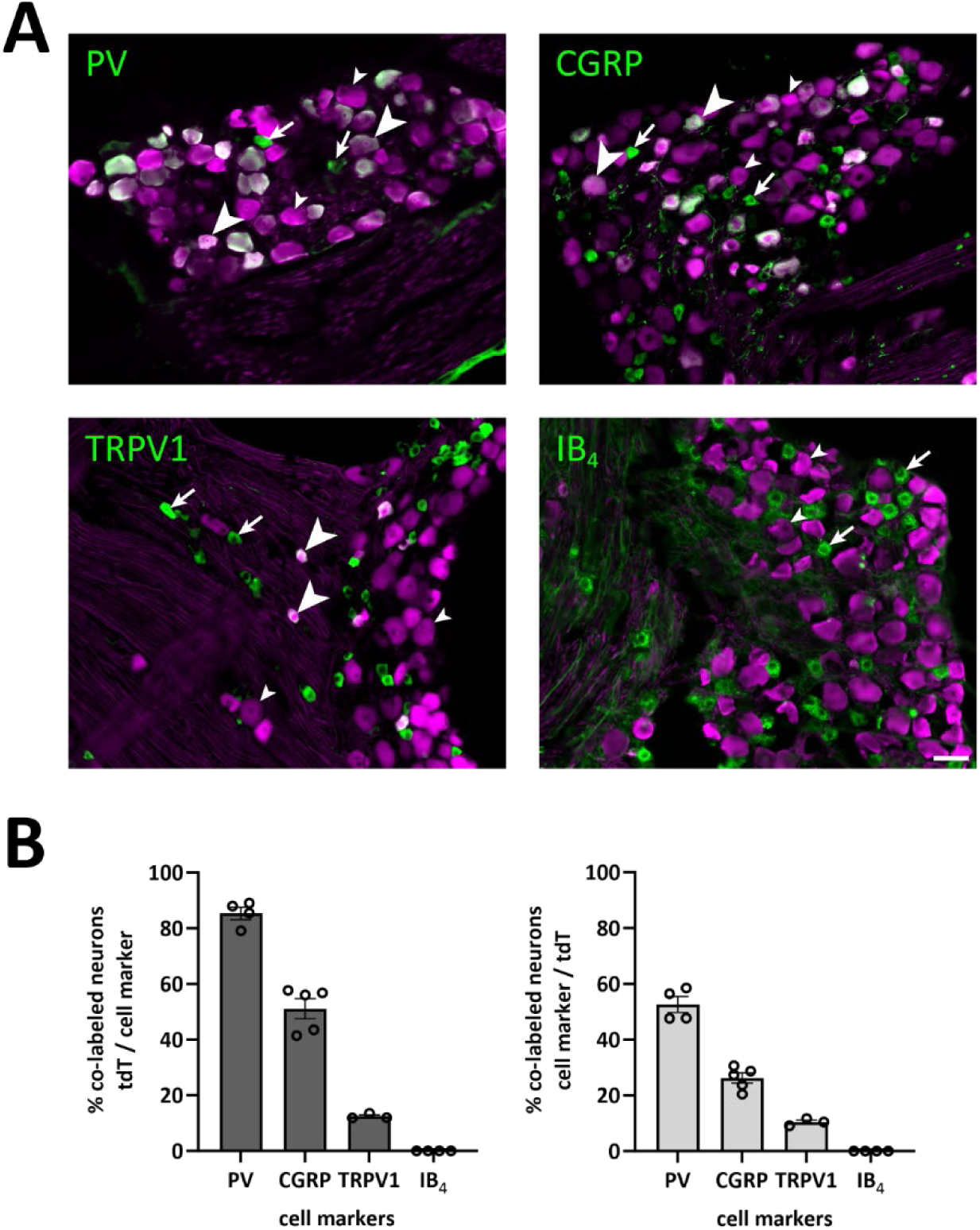
Co-localization with markers of neuronal populations in the DRG. **A**, DRG sections labeled for tdTomato and parvalbumin (PV), CGRP, TRPV1 or IB_4_ binding**. Arrows indicate cells immunoreactive for cell marker but not tdTomato; small arrowheads indicate tdTomato^+^ cells not immunoreactive for the cell marker; large arrowheads indicate cells immunoreactive for both tdTomato and cell marker. B**, quantification of co-localization between tdTomato^+^ cells and the respective markers. Each data point represents a single section and DRG; n=2 or 3 mice per marker.

In spinal cord white matter, tdTomato^+^ fibers were numerous in the dorsal column, in accordance with the extensive projection of myelinated PAFs to dorsal column nuclei (Fig. 5A). The corticospinal tract in the ventral dorsal funiculus showed scattered tdTomato^+^ fibers, in line with the recombination observed in some cortical layer 5 neurons (see below). Occasional tdTomato^+^ fibers were also found in the ventrolateral funiculus of the spinal cord. In gray matter, motor neurons and some neurons of the lateral spinal nucleus exhibited tdTomato expression, but elsewhere in the spinal gray matter tdTomato^+^ cell bodies were very infrequent. tdTomato^+^ processes were predominantly found in the dorsal horn, as well as in lamina IX and to a lesser extent in lamina VII; passing fibers were evident in other laminae. In the superficial dorsal horn, tdTomato^+^ processes were mainly found in lamina I and extending into the dorsal part of outer lamina II, essentially overlapping with the pattern of CGRP-IR and TRPV1-IR fibers (Fig. 5B), and in inner lamina II ventral to the IB_4_ binding band, overlapping with the main band of PKCγ-IR neurons (Fig. 5C); thus the central portion of lamina II, which in the mouse is occupied by IB_4_ binding non-peptidergic C fiber terminals, was essentially devoid of tdTomato^+^ fibers. The deep dorsal horn was densely populated by tdTomato^+^ processes in laminae III-IV and in medial lamina V and VI, but less so in lateral lamina V. In the brain, only scattered neurons showed tdTomato expression after one tamoxifen injection (Fig. 5D), and among identified populations generally only a proportion of cells showed recombination. We did not perform a comprehensive survey of recombined cells; however, in the cerebellum a relatively large fraction of Purkinje cells exhibited tdTomato, as did mossy fiber terminals in the granule cell layer. In the neocortex a small portion of layer 5 pyramidal neurons were tdTomato^+^, as were some cells in the thalamus. As expected, neurons of the trigeminal motor nucleus were tdTomato^+^, as were numerous processes in the spinal trigeminal nucleus.

**Figure 5.**
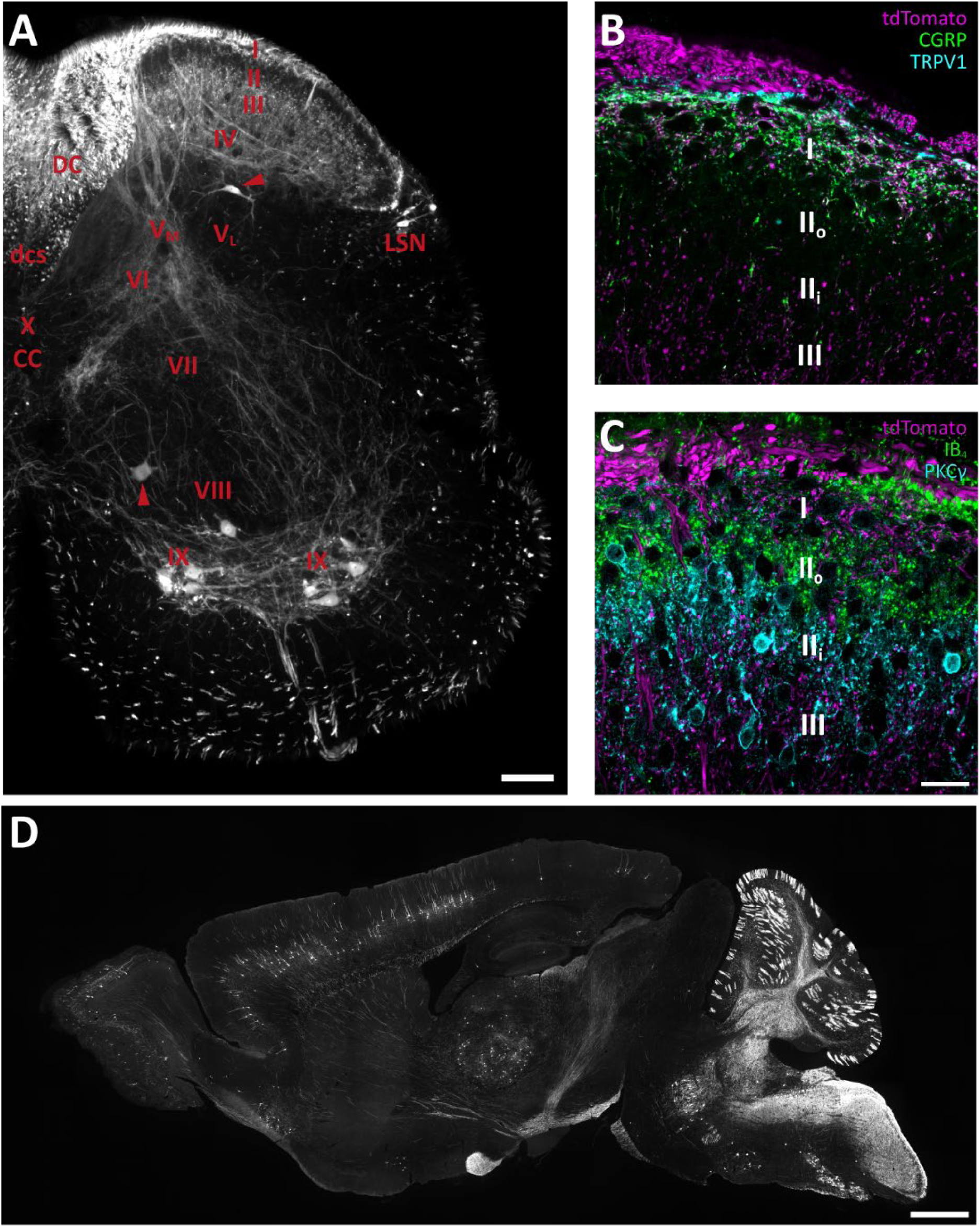
Reporter-expressing processes in the spinal cord and brain. **A**, distribution of tdTomato^+^ processes and cells in a transverse hemisection of spinal cord at the L4 level. Roman numerals denote Rexed’s laminae; V_M_ and V_L_ indicate medial and lateral lamina V, respectively. Arrowheads indicate tdTomato^+^ cell bodies outside lamina IX. *CC*, central canal; *DC*, dorsal column; *dcs*, dorsal corticospinal tract; *LSN*, lateral spinal nucleus. **B**, higher-magnification view of the superficial dorsal horn in a section co-labeled for tdTomato, CGRP and TRPV1. Roman numerals indicate Rexed’s laminae; II_o_ and II_i_ indicate outer and inner lamina II, respectively. **C**, as B but labeled for PKCγ and IB_4_ binding sites. **D**, a parasagittal brain section showing tdTomato^+^ cells and processes. Scale bar in A, 100 µm. Scale bar in C, 20 µm, valid for B and C. Scale bar in D, 1 mm.

To confirm the utility of the *Nefh*^CreERT2^ mouse for morphological and physiological studies at different scales, we first crossed the mouse line with an LSL-APEX2 mouse line, which expresses the peroxidase APEX2 in the mitochondrial matrix in a Cre-dependent manner [19,26], allowing investigation of the ultrastructure of myelinated PAFs in the spinal cord by electron microscopy. In spinal cord tissue from these mice we noted widespread peroxidase reaction product throughout the dorsal horn; in the electron microscope, numerous nerve endings harboring mitochondria with peroxidase reaction product were found throughout the dorsal horn, with the exception of dorsal lamina II, were such terminals were scarce (Fig. 6). Many of these nerve endings formed complex synaptic glomeruli, as is characteristic of many primary afferent fibers [27,28]. In the ventral horn, motor neuron somata and dendrites possessed mitochondria with peroxidase reaction product, as did nerve terminals of presumed primary afferent origin in this region. Notably, in lamina IX we observed terminals with peroxidase reactive mitochondria contacting dendrites that also had mitochondria with peroxidase reaction product; these synapses are likely connections between type Ia fibers and motor neuron dendrites.

**Figure 6.**
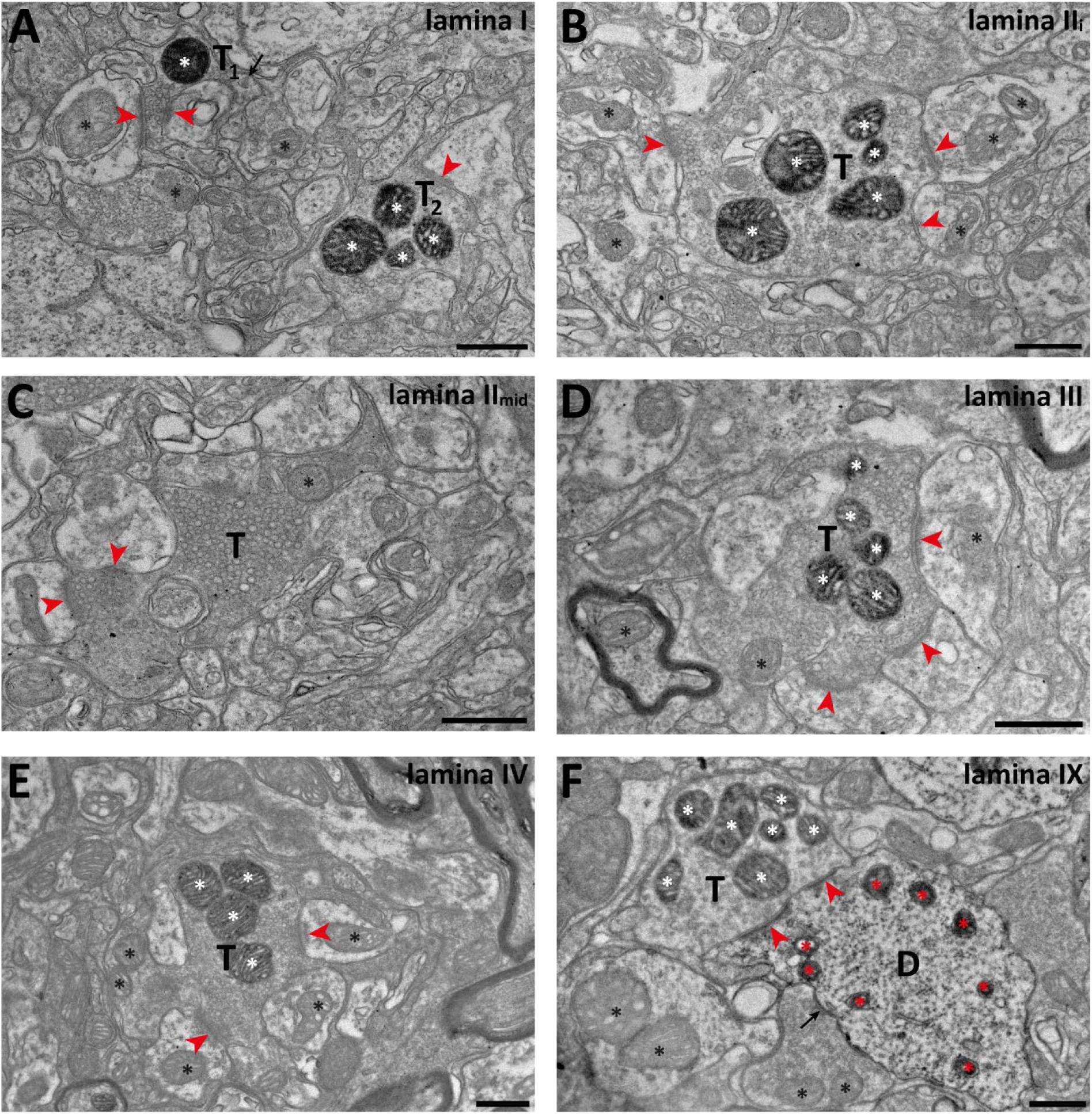
Electron microscopy of reporter-expressing nerve endings in the spinal dorsal horn. **A**, two putative myelinated nociceptor terminals (*T*_1_ and *T*_2_) with peroxidase reactive mitochondria (white asterisks) in lamina I of the dorsal horn. Synapses formed by these terminals are indicated by arrowheads. The synapses formed by T_1_ may be a single saddle synapse. Black asterisks indicate examples of non-labeled mitochondria. Arrow indicates a dense core vesicle in T_1_. **B**, a peroxidase reactive glomerular terminal (*T*) in inner lamina II (II_i_); symbols are used as in A. **C**, a type I glomerulus in mid-third lamina II. Note that the mitochondrion (asterisk) in the central C fiber terminal (*T*) does not contain peroxidase reaction product. **D** and **E**, Glomerular terminals (*T*) in lamina III and IV respectively with peroxidase reactive mitochondria. **F**, a putative type Ia fiber terminal (*T*) with reactive mitochondria (white asterisks) forming a perforated synapse (arrowheads) with a putative motor neuron dendrite (*D*) also harboring peroxidase reactive mitochondria (red asterisks). A terminal with non-reactive mitochondria also forms a synapse (arrow) with the same dendrite. Scale bars are 500 nm in all panels.

Finally, we crossed *Nefh*^CreERT2^ mice with RCL-GCaMP6f mice to enable measurement of their physiological response *in vivo* through Ca^2+^ imaging of DRG neurons (Fig. 7) [22]. Since multiple myelinated classes have been shown to be enriched in the DRGs of the sacral nerve [29,30], we chose to record neurons in the S1 ganglion. As expected, in these ganglia, GCaMP6f expressing neurons were readily activated by a variety of low-threshold mechanical stimuli (Fig. 7B) whereas cells responding to temperature stimuli were very scarce, probably reflecting very high threshold myelinated polymodal nociceptors (Fig. 7C). Next, we wished to test whether LTMRs were enriched among recombined neurons in the *Nefh*^CreERT2^;GCaMP6f mouse, and recorded responses to brushing, hair pull and pinch in these mice and in pan-GCaMP6f mice where neurons within all sensory classes expressed GCaMP6f (Fig. 7D,E) (Szczot et al. 2018). As expected, LTMRs were a significantly larger fraction of responders in *Nefh*^CreERT2^;GCaMP6f mice as compared to the pan-GCaMP6f mice, where also C fibers expressed GCaMP6f.

**Figure 7.**
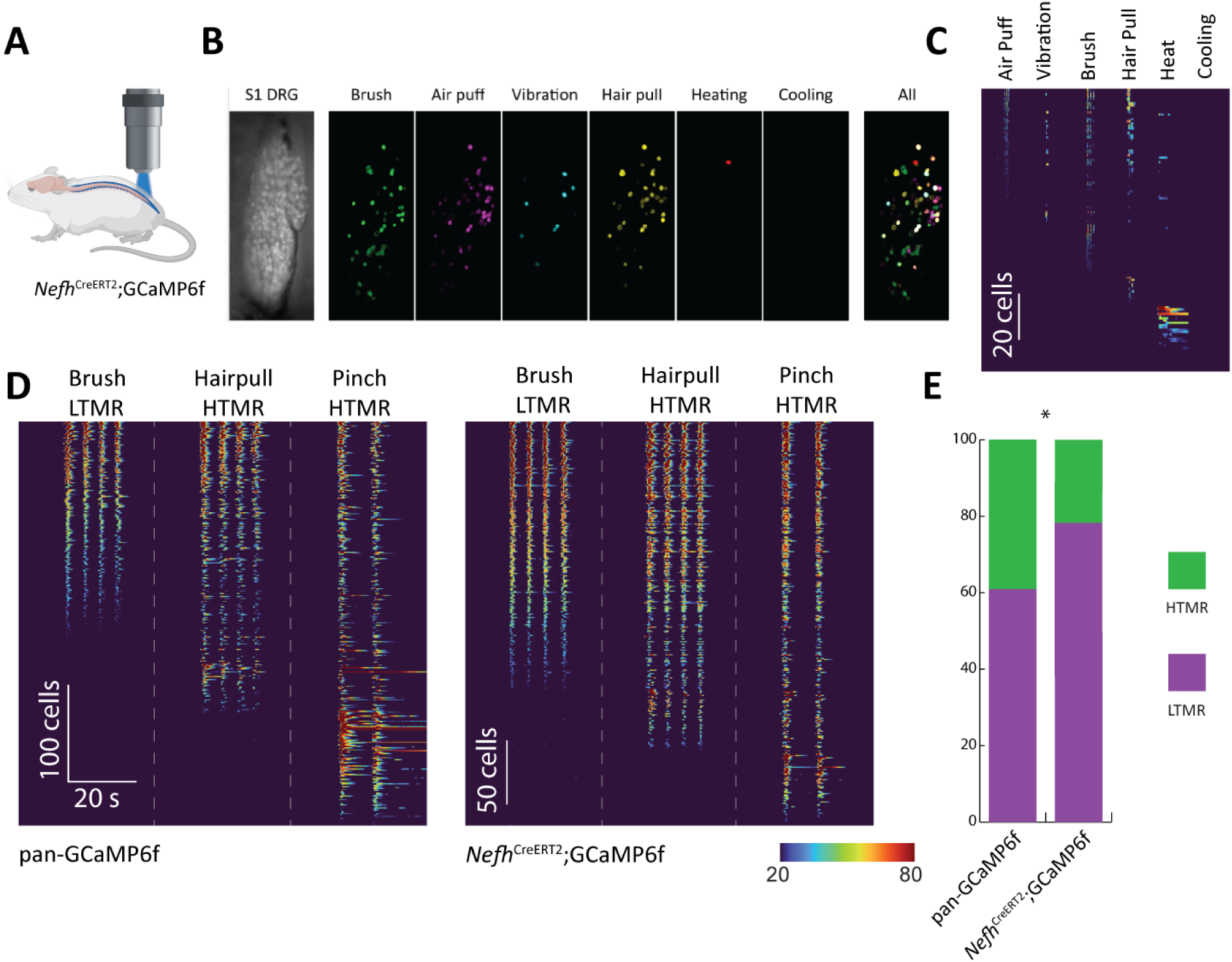
*Nefh*^CreRT2^ mice allows robust targeting of myelinated primary sensory neurons for physiological characterization. **A**, Schematic of *Nefh*^CreERT2^ driver mice crossed with conditional RCL-GCaMP6f reporter mice (*Nefh*^CreERT2^;GCaMP6f); this cross was used for *in vivo* Ca^2+^ imaging of the activity of myelinated DRG neurons. **B**, *Nefh*^CreERT2^ targets multiple classes of specialized LTMRs and HTMRs. Leftmost panel, baseline fluorescence of ganglion surface; center panels, projection visualizing neurons activated by indicated stimuli; rightmost panel pseudocolored overlay showing combinatorial activity. **C**, Quantitative heatmap showing differential responses of cells to the indicated stimuli. **D**, Comparison of LTMR and HTMR cells activated in *Nefh*^CreERT2^;GCaMP6f line (n=4 mice, 313 cells) and in pan-neuronal GCaMP6f-expressing mice (pan-GCaMP6f; n=4 mice, 362 cells). **E,** Proportion of LTMRs among activated sacral neurons in *Nefh*^CreERT2^;GCaMP6f mice is significantly higher than in sacral ganglia of pan-GCaMP6f mice (Fisher’s exact test, *p* < 0.0001).

## Discussion

Here we generated and characterized a knock-in *Nefh^CreERT2^* mouse line, showing that this line is a versatile and selective tool for targeting myelinated PAFs to the exclusion of unmyelinated C fibers. One injection of 2 mg tamoxifen in juvenile mice proved sufficient for highly efficient (>85 %) and selective (>95 %) recombination in NFH^+^ DRG neurons, while a single injection of 20 µg tamoxifen was shown to be useful for sparse labeling of such neurons. However, even with five tamoxifen injections, recombination in CNS neurons remained rather restricted. Nevertheless, neurons that did show recombination were distributed in accordance with *Nefh* mRNA [31,32]. Previously reported transgenic mouse lines constitutively expressing Cre under the *Nefh* promoter via random integration shows more widespread recombination in the CNS, whereas DRG expression was not evaluated in these lines [33].

For morphological and functional characterization of myelinated PAFs and to assess the utility of our newly generated *Nefh^CreERT2^* mice, we crossed these mice with several different reporter mouse lines, including Ai14 (LSL-tdTomato) mice for light microscopy, LSL-APEX2 mice for electron microscopy and RCL-GCaMP6f mice for *in vivo* Ca^2+^ imaging in DRGs. Observations obtained from these mice were in close agreement with known characteristics of the broad population of myelinated PAFs. In L3-L5 DRGs of *Nefh^CreERT2^*;Ai14 mice, neurons expressing tdTomato were medium-to-large sized, and co-expression with neurochemical markers confirmed recombination in essentially all myelinated DRG neurons, including proprioceptors and myelinated peptidergic nociceptors [15,24,34]. In the skin of these mice, tdTomato^+^ fibers were found to form circumferential and lanceolate endings around hair follicles, as well as endings associated with Meissner corpuscles. Free epidermal nerve endings, likely belonging to myelinated nociceptors, were also observed. In the spinal cord, motor neurons showed recombination whereas few neurons did elsewhere in the gray matter, including in the dorsal horn. Axons containing tdTomato were most dense in the dorsal columns and in lamina I and laminae III-IV, whereas projections of presumed proprioceptive origin were found as expected in intermediate and ventral laminae. At the electron microscopic level, nerve terminals showing recombination-dependent peroxidase labeling had morphological characteristics in line with previous observations of myelinated primary afferent terminals, while terminals that originate from non-peptidergic C fiber nociceptors did not have reactive mitochondria [25,27,35,36].

Notably, in the ventral horn we encountered synaptic connections where both the presynaptic terminal and the postsynaptic dendrite had peroxidase-containing mitochondria. Since type Ia fibers are the only primary afferent fibers that monosynaptically contact motor neurons, and corticospinal and intraspinal pathways show only sparse recombination in these mice, it is likely that many of these connections are type Ia-motor neuron synapses. *In vivo* Ca^2+^ imaging in S1 ganglia mice showed the expected patterns of cellular responses to cutaneous stimulation – many cells were activated by one or more of low-threshold mechanical stimuli such as air puffs, vibration or gentle brushing, whereas a small proportion of cells were activated exclusively by hair pull [22,37,38]. In addition, LTMRs represented significantly larger fraction of responding cells than in the general population captured in pan-GCaMP6f mice. In conclusion, these observations indicate that the *Nefh^CreERT2^* mouse line is able to, with very high specificity and selectivity, target myelinated DRG neurons.

**Whereas recombination in the DRG occurred selectively in myelinated neurons, it should be noted that there was widespread, although in most regions relatively sparse, recombination also in the CNS. Therefore, if the experimental design necessitates exclusive targeting of DRG neurons, an intersectional targeting approach may be employed, where the combined expression of Nefh and the second marker gene is restricted to DRGs.**

Although the functional significance of the distinction between myelinated and unmyelinated sensory neurons was apparent by the 1930’s [7,8], and despite the rapid increase in the use of genetic strategies for the visualization and manipulation of sensory systems, no genetic tools have been available to reliably distinguish between these two neuronal populations. Here we describe a new driver mouse line that can be used to robustly and selectively capture myelinated primary afferent neurons in intersectional genetic targeting strategies. Thus we expect that this mouse will be a valuable addition to the toolbox used for functional and anatomical characterization of subpopulations of somatosensory neurons.

## Acknowledgments

This study was funded by Knut and Alice Wallenberg Foundation, project no. 2019.0047. We thank Dr. Maria Ntzouni and Dr. Vesa Loitto at the Microscopy Core Facility at the Medical Faculty, Linköping University for technical assistance, and Olga Gewartowska and Jakub Gruchota at the International Institute of Molecular and Cell Biology, Warsaw, Poland for optimizing the CRISPR targeting and zygote injections.

## Author contributions

M.L. conceived the project; J.C.Y.C., L.K., F.M.F., M.S., W.S.J. and M.L. designed experiments; J.C.Y.C., L.K., F.M.F., and M.L. performed experiments and analyzed data; M.L. wrote the paper with feedback from all authors.

## Data availability

Data collected in this study is available from the corresponding author upon reasonable request.

## Additional information

The authors declare no competing interests.

## References

1 Usoskin, D. et al. Unbiased classification of sensory neuron types by large-scale single-cell rna sequencing. Nat. Neurosci. 18, 145 (2015). 10.1038/nn.3881

2 Kupari, J. et al. Single cell transcriptomics of primate sensory neurons identifies cell types associated with chronic pain. Nat Commun 12, 1510 (2021). 10.1038/s41467-021-21725-z

3 Sharma, N. et al. The emergence of transcriptional identity in somatosensory neurons. Nature 577, 392–398 (2020). 10.1038/s41586-019-1900-1

4 Li, C.-L. et al. Somatosensory neuron types identified by high-coverage single-cell rna-sequencing and functional heterogeneity. Cell Res. 26, 83 (2015). 10.1038/cr.2015.149

5 Yu, H. et al. Leveraging deep single-soma rna sequencing to explore the neural basis of human somatosensation. Nat. Neurosci. (2024). 10.1038/s41593-024-01794-1

6 Zeisel, A. et al. Molecular architecture of the mouse nervous system. Cell 174, 999–1014.e1022 (2018). 10.1016/j.cell.2018.06.021

7 Perl, E. R. Ideas about pain, a historical view. Nat. Rev. Neurosci. 8, 71–80 (2007). 10.1038/nrn2042

8 Willis, W. D. & Coggeshall, R. E. Sensory mechanisms of the spinal cord. 3rd edn, (Kluwer, 2004).

9 Olausson, H. et al. Unmyelinated tactile afferents signal touch and project to insular cortex. Nat Neurosci 5, 900–904 (2002). 10.1038/nn896

10 Löken, L. S., Wessberg, J., Morrison, I., McGlone, F. & Olausson, H. Coding of pleasant touch by unmyelinated afferents in humans. Nat. Neurosci. 12, 547 (2009). 10.1038/nn.2312

11 Zotterman, Y. Touch, pain and tickling: An electro-physiological investigation on cutaneous sensory nerves. J. Physiol. 95, 1–28 (1939).

12 Nagi, S. S. et al. An ultrafast system for signaling mechanical pain in human skin. Sci Adv 5, eaaw1297 (2019). 10.1126/sciadv.aaw1297

13 Djouhri, L. & Lawson, S. N. Aβ-fiber nociceptive primary afferent neurons: A review of incidence and properties in relation to other afferent a-fiber neurons in mammals. Brain Res. Rev. 46, 131–145 (2004).

14 Lawson, S. N., Harper, A. A., Harper, E. I., Garson, J. A. & Anderton, B. H. A monoclonal antibody against neurofilament protein specifically labels a subpopulation of rat sensory neurones. J Comp Neurol 228, 263–272 (1984). 10.1002/cne.902280211

15 Lawson, S. N. & Waddell, P. J. Soma neurofilament immunoreactivity is related to cell size and fibre conduction velocity in rat primary sensory neurons. J. Physiol. 435, 41–63 (1991). 10.1113/jphysiol.1991.sp018497

16 Kaczmarczyk, L. et al. Slc1a3-2a-creert2 mice reveal unique features of bergmann glia and augment a growing collection of cre drivers and effectors in the 129s4 genetic background. Scientific reports 11, 5412 (2021). 10.1038/s41598-021-84887-2

17 Madisen, L. et al. A robust and high-throughput cre reporting and characterization system for the whole mouse brain. Nat. Neurosci. 13, 133–140 (2010). 10.1038/nn.2467

18 Madisen, L. et al. Transgenic mice for intersectional targeting of neural sensors and effectors with high specificity and performance. Neuron 85, 942–958 (2015). 10.1016/j.neuron.2015.02.022

19 Zhang, Q., Lee, W.-C. A., Paul, D. L. & Ginty, D. D. Multiplexed peroxidase-based electron microscopy labeling enables simultaneous visualization of multiple cell types. Nat. Neurosci. 22, 828–839 (2019). 10.1038/s41593-019-0358-7

20 Raymond, C. S. & Soriano, P. High-efficiency flp and φc31 site-specific recombination in mammalian cells. PloS one 2, e162 (2007). 10.1371/journal.pone.0000162

21 Szczot, M. et al. Piezo2 mediates injury-induced tactile pain in mice and humans. Science Translational Medicine 10, eaat9892 (2018). doi:10.1126/scitranslmed.aat9892

22 Ghitani, N. et al. Specialized mechanosensory nociceptors mediating rapid responses to hair pull. Neuron 95, 944–954.e944 (2017). 10.1016/j.neuron.2017.07.024

23 Shaw, G. & Weber, K. Differential expression of neurofilament triplet proteins in brain development. Nature 298, 277–279 (1982). 10.1038/298277a0

24 Arber, S., Ladle, D. R., Lin, J. H., Frank, E. & Jessell, T. M. Ets gene er81 controls the formation of functional connections between group ia sensory afferents and motor neurons. Cell 101, 485–498 (2000). 10.1016/S0092-8674(00)80859-4

25 Gerke, M. B. & Plenderleith, M. B. Ultrastructural analysis of the central terminals of primary sensory neurones labelled by transganglionic transport of bandeiraea simplicifolia i-isolectin b4. Neuroscience 127, 165–175 (2004).

26 Lam, S. S. et al. Directed evolution of apex2 for electron microscopy and proximity labeling. Nat Methods 12, 51–54 (2015). 10.1038/nmeth.3179

27 Maxwell, D. J. & Réthelyi, M. Ultrastructure and synaptic connections of cutaneous afferent fibres in the spinal cord. Trends Neurosci 10, 117–123 (1987). 10.1016/0166-2236(87)90056-7

28 Ribeiro-da-Silva, A. in The rat nervous system (ed G. Paxinos) 129–148 (Academic Press, 2004).

29 Lam, R. M. et al. Piezo2 and perineal mechanosensation are essential for sexual function. Science 381, 906–910 (2023). doi:10.1126/science.adg0144

30 Qi, L. et al. Krause corpuscles are genital vibrotactile sensors for sexual behaviours. Nature 630, 926–934 (2024). 10.1038/s41586-024-07528-4

31 Roussel, G. et al. In situ localization of nf-h neurofilament subunit mrnas in rat brain. Developmental Neuroscience 13, 98–103 (1991). 10.1159/000112146

32 Allen mouse brain atlas [dataset]. http://mouse.brain-map.org (2011).

33 Hirasawa, M. et al. Neuron-specific expression of cre recombinase during the late phase of brain development. Neurosci. Res. 40, 125–132 (2001). 10.1016/S0168-0102(01)00216-4

34 Lawson, S. N., Crepps, B. & Perl, E. R. Calcitonin gene-related peptide immunoreactivity and afferent receptive properties of dorsal root ganglion neurones in guinea-pigs. The Journal of Physiology 540, 989–1002 (2002). 10.1113/jphysiol.2001.013086

35 Larsson, M. & Broman, J. Synaptic organization of vglut3 expressing low-threshold mechanosensitive c fiber terminals in the rodent spinal cord. eNeuro, ENEURO.0007-0019.2019 (2019). 10.1523/eneuro.0007-19.2019

36 Light, A. R. & Perl, E. R. Spinal termination of functionally identified primary afferent neurons with slowly conducting myelinated fibers. J Comp Neurol 186, 133–150 (1979). 10.1002/cne.901860203

37 von Buchholtz, L. J. et al. Decoding cellular mechanisms for mechanosensory discrimination. Neuron 109, 285–298.e285 (2021). 10.1016/j.neuron.2020.10.028

38 Bouchatta, O. et al. Piezo2-dependent rapid pain system in humans and mice. bioRxiv, 2023.2012.2001.569650 (2023). 10.1101/2023.12.01.569650

