## Supplementary Fig. 1 for "Genetic targeting of myelinated primary afferent neurons using a new *Nefh^CreERT2^* knock-in mouse"

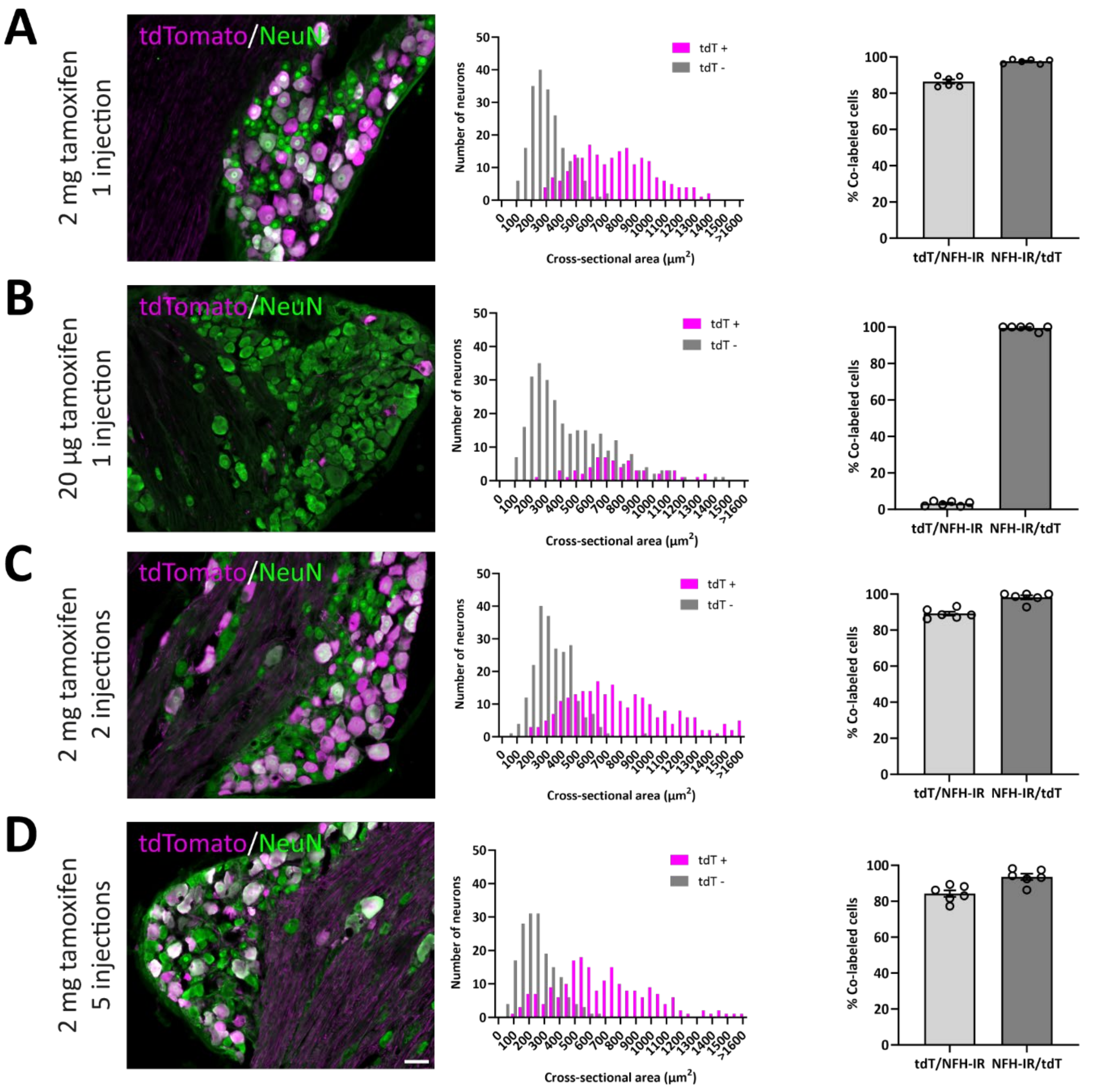


**Figure S1.** Soma size and recombination efficiency in lumbar DRGs after different tamoxifen administration regimes. **A**, a single low-dose (20 µg) tamoxifen injection. **B-D**, one (B), two (C) or five (D) injections of 2 mg tamoxifen. Left panels, representative images of DRGs showing tdTomato expression among NeuN immunoreactive neurons. Middle panels, size distributions of tdTomato$^+$ and tdTomato$^-$ DRG neurons. Right panels, recombination efficiency (mean ± S.E.M). with respect to NFH-IR. n=3 mice.  Scale bar in D is 50 µm, valid for all panels.
